# Link Between Short tandem Repeats and Translation Initiation Site Selection

**DOI:** 10.1101/316950

**Authors:** M Arabfard, K Kavousi, A Delbari, M Ohadi

## Abstract

Recent work in yeast and humans suggest that evolutionary divergence in *cis*-regulatory sequences impact translation initiation sites (TISs). *Cis*-elements can also affect the efficacy and amount of protein synthesis. Despite their vast biological implication, the landscape and relevance of short tandem repeats (STRs)/microsatellites to the human protein-coding gene TISs remain largely unknown. Here we characterized the STR distribution at the 120 bp cDNA sequence upstream of all annotated human protein-coding gene TISs based on the Ensembl database. Furthermore, we performed a comparative genomics study of all annotated orthologous TIS-flanking sequences across 47 vertebrate species (755,956 transcripts), aimed at identifying human-specific STRs in this interval. We also hypothesized that STRs may be used as genetic codes for the initiation of translation. The initial five amino acid sequences (excluding the initial methionine) that were flanked by STRs in human were BLASTed against the initial orthologous five amino acids in other vertebrate species (2,025,817 pair-wise TIS comparisons) in order to compare the number of events in which human-specific and non-specific STRs occurred with homologous and non-homologous TISs (i.e. ≥50% and <50% similarity of the five amino acids). We characterized human-specific STRs and a bias of this compartment in comparison to the overall (human-specific and non-specific) distribution of STRs (Mann Whitney p=1.4 × 10^−11^). We also found significant enrichment of non-homologous TISs flanked by human-specific STRs (p<0.00001). In conclusion, our data indicate a link between STRs and TIS selection, which is supported by differential evolution of the human-specific STRs in the TIS upstream flanking sequence.

**Abbreviations:**

cDNA: Complementary DNA
CDS: Coding DNA sequence
STR: Short Tandem Repeat
TIS: Translation Initiation Site
TSS: Transcription Start Site

## Introduction

An increasing number of human protein-coding genes are unraveled to consist of alternative translation initiation sites (TISs), which are selected based on complex and yet not fully known scanning mechanisms (Andreev et al. 2017; Lee et al. 2012). The alternative TISs result in various protein structures and functions (Fukushima et al. 2012; Georgii et al. 2011). Selection of TISs and the level of translation and protein synthesis depend on the *cis* regulatory elements in the mRNA sequence and its secondary structure such as the formation of hair-pins and thermal stability (Cenik et al; Babendure et al. 2006; Master et al. 2016).

One of the important and understudied *cis*-regulatory elements affecting translation are short tandem repeats (STRs)/microsatellites. In physiological terms, STRs can dramatically influence TIS and the amount of protein synthesis. Poly(A) tracts in the 5′- untranslated region (UTR) are important sites for translation regulation in yeast. These poly(A) tracts can interact with translation initiation factors or poly(A) binding proteins (PABP) to either increase or decrease translation efficiency. Pre-AUG A_N_ can enhance internal ribosomal entry both in the presence of PABP and eIF-4G in *Saccharomyces cerevisiae* (Gilbert *et al.* 2007), and in the complete absence of PABP and eIF-4G (Shirokikh and Spirin 2008). Biased distribution of dinucleotide repeats is a known phenomenon in the region immediately upstream of the TISs in *E. coli* (Yamagishi et al. 2002). In pathological instances, expansion of STRs in the RNA structure result in toxic RNAs and non-AUG translation, and the development of several human-specific neurological disorders (Glineburg et al. 2018; Rovozzo et al. 2016; Krauss et al. 2013).

Genome-scale findings of the evolutionary trend of a number of STRs has begun to unfold their implications in respect with speciation and species-specific characteristics/phenotypes (Yuan et al. 2018; Emamalizadeh et al. 2017; Abe and Gemmell 2016; Bushehri et al. 2016; Namdar-Aligoodarzi et al. 2016; Nikkhah et al. 2016; Bilgin Sonay et al. 2015; Rezazadeh et al. 2014;Khademi et al. 2017; Mohammadparast et al. 2014; Ohadi et al. 2012; King et al. 2012). The hypermutable nature of STRs and their large unascertained reservoir of functionality make them an ideal source of evolutionary adaptation, speciation, and disease (Hannan et al. 2018;Bagshaw et al. 2017; Press et al. 2017; Ohadi et al.2015; Valipour et al. 2013; Heidari et al. 2012). In line with the above, recent reports indicate a role of repetitive sequences in the creation of new transcription start sites (TSSs) in human (Nazaripanah et al. 2018; Alizadeh et al. 2018; Li et al. 2018; Kramer et al. 2013).

Here, we performed a genome-scale screen of the upstream complementary DNA (cDNA) sequence flanking all human protein-coding TISs annotated in the Ensembl database and a comparative genomics analysis to examine a possible link between these STRs and TIS selection in 47 species encompassing the major classes of vertebrates.

## Results

### Genome-scale Distribution of STRs in the 120 bp upstream sequence of TISs in human

Mono and dinucleotide STRs dominated STRs of >1000 counts, and the (T)6 mononucleotide repeat was the most abundant STR in this interval, succeeded by the (CT)3 and (TC)3 dinucleotide STRs (Fig. 1).

**Fig. 1.**
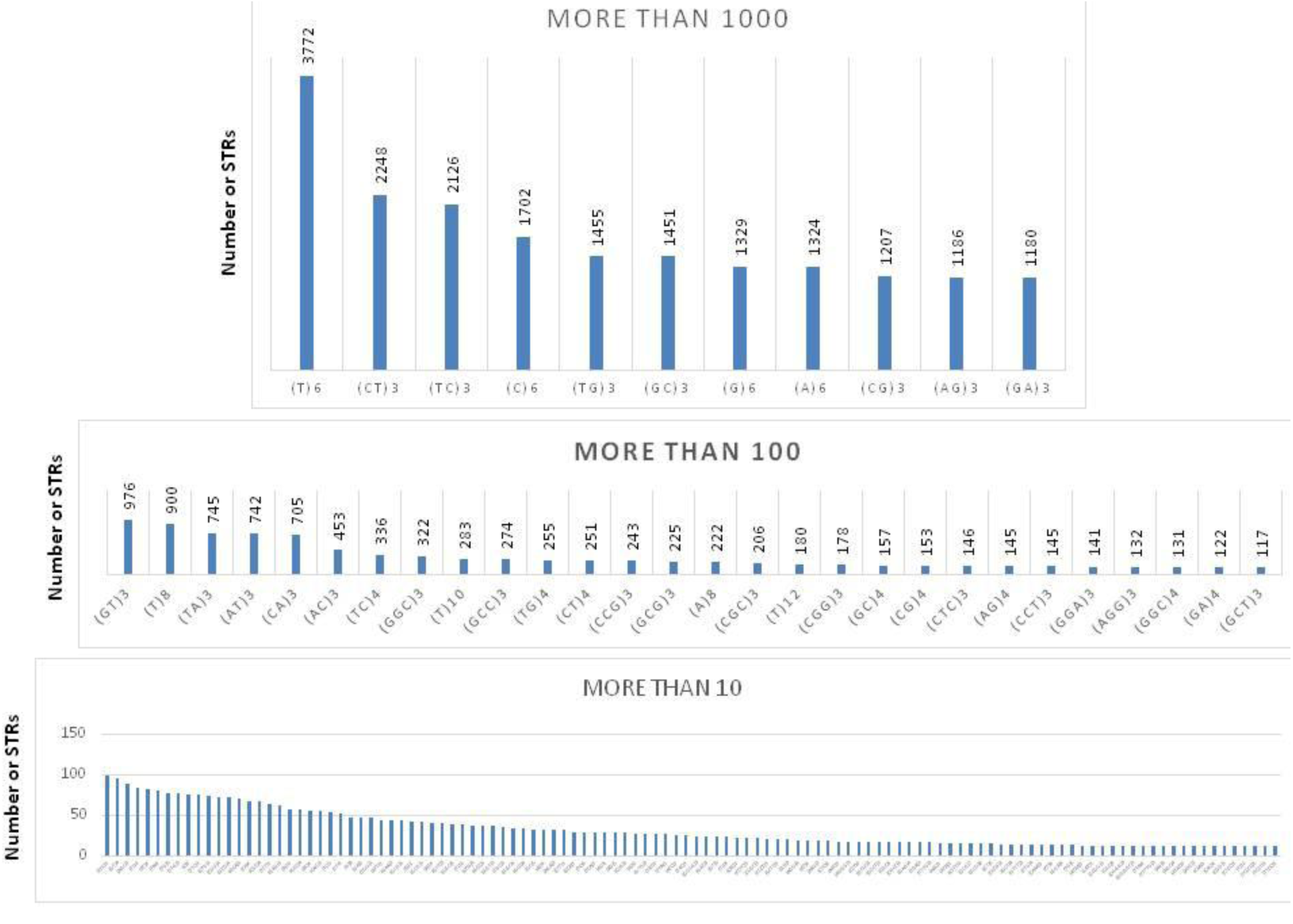
Genome-scale landscape of the human STRs in the 120 bp cDNA sequence flanking TISs. The abundance of STRs is sorted in the ascending order.

Trinucleotide STRs were less abundant, observed at counts between 100 and 1000, and predominated by GC-rich composition such as (GGC)3, (GCC)3, (CCG)3 and (GCG)3. In the non-GC composition, (CTC)3 and (CCT)3 were the most common trinucleotide STRs. Tetra, penta, and hexanucleotide STRs were at lesser abundance than the above categories and observed at <100 counts, where (GGCG)3 and (CCTC)3 were the most abundant tetranucleotide STRs. Only three pentanucleotide STR classes, (GGGGC)3, (TTTTG)3 and (TCCCC)3 were observed at counts >10 in the screened interval.

### Human-specific STR fingerprints in the TIS-flanking sequence and skewing of this compartment in comparison to the overall human STR distribution

Two thousand two hundred sixty eight genes contained human-specific TIS-flanking STRs, which were of a wide range of nucleotide compositions of mono, di, tri, tetra, penta, and hexanucleotide repeats (Table 1).

**Table 1.** The 5^th^ percentile of human protein-coding genes which contain human-specific STRs in their TIS-flanking sequence.

| Gene Name | Gene Ensembl ID | Gene Transcript ID | STR |
| --- | --- | --- | --- |
| ***NVL*** | ENSG00000143748 | ENST00000436927 | (T)22 |
| ***ELOF1*** | ENSG00000130165 | ENST00000587806 | (T)20 |
| ***OR4K2*** | ENSG00000165762 | ENST00000641885 | (T)20 |
| ***LRRC19*** | ENSG00000184434 | ENST00000380055 | (A)20 |
| ***MGRN1*** | ENSG00000102858 | ENST00000591895 | (A)18 |
| ***MGRN1*** | ENSG00000102858 | ENST00000591895 | (A)18 |
| ***SULT1A3*** | ENSG00000261052 | ENST00000338971 | (A)18 |
| ***SULT1A3*** | ENSG00000261052 | ENST00000395138 | (A)18 |
| ***TBR1*** | ENSG00000136535 | ENST00000410035 | (TG)17 |
| ***RORA*** | ENSG00000069667 | ENST00000335670 | (T)16 |
| ***RORA*** | ENSG00000069667 | ENST00000335670 | (T)16 |
| ***ADAP2*** | ENSG00000184060 | ENST00000581548 | (A)16 |
| ***DDX20*** | ENSG00000064703 | ENST00000475700 | (A)16 |
| **GDI2** | ENSG00000057608 | ENST00000380127 | (T)16 |
| **GDI2** | ENSG00000057608 | ENST00000609712 | (T)16 |
| **GSTA4** | ENSG00000170899 | ENST00000370960 | (A)16 |
| **SULT1A4** | ENSG00000213648 | ENST00000360423 | (A)16 |
| **ZNF283** | ENSG00000167637 | ENST00000618787 | (T)16 |
| **ZNF283** | ENSG00000167637 | ENST00000593268 | (T)16 |
| **SGIP1** | ENSG00000118473 | ENST00000435165 | (A)16 |
| **OR1C1** | ENSG00000221888 | ENST00000641256 | (CA)15 |
| **CNGA1** | ENSG00000198515 | ENST00000402813 | (CA)15 |
| **POLR2F** | ENSG00000100142 | ENST00000492213 | (T)14 |
| **SNX19** | ENSG00000120451 | ENST00000528555 | (T)14 |
| **SNX19** | ENSG00000120451 | ENST00000530356 | (T)14 |
| **OR7A10** | ENSG00000127515 | ENST00000641129 | (CT)14 |
| **HIGD2B** | ENSG00000175202 | ENST00000311755 | (A)14 |
| **LCA5L** | ENSG00000157578 | ENST00000288350 | (A)14 |
| **LCA5L** | ENSG00000157578 | ENST00000358268 | (A)14 |
| **LCA5L** | ENSG00000157578 | ENST00000485895 | (A)14 |
| **LCA5L** | ENSG00000157578 | ENST00000418018 | (A)14 |
| **LCA5L** | ENSG00000157578 | ENST00000448288 | (A)14 |
| **LCA5L** | ENSG00000157578 | ENST00000434281 | (A)14 |
| **LCA5L** | ENSG00000157578 | ENST00000438404 | (A)14 |
| **LCA5L** | ENSG00000157578 | ENST00000411566 | (A)14 |
| **LCA5L** | ENSG00000157578 | ENST00000415863 | (A)14 |
| **LCA5L** | ENSG00000157578 | ENST00000426783 | (A)14 |
| **LCA5L** | ENSG00000157578 | ENST00000456017 | (A)14 |
| **LRRC36** | ENSG00000159708 | ENST00000569499 | (T)14 |
| **LRRC36** | ENSG00000159708 | ENST00000568804 | (T)14 |
| **MRPL13** | ENSG00000172172 | ENST00000518918 | (T)14 |
| **TEX11** | ENSG00000120498 | ENST00000395889 | (TTCC)14 |
| **GALK2** | ENSG00000156958 | ENST00000560654 | (TG)13 |
| **GALK2** | ENSG00000156958 | ENST00000396509 | (TG)13 |
| **GALK2** | ENSG00000156958 | ENST00000558145 | (TG)13 |
| **GALK2** | ENSG00000156958 | ENST00000544523 | (TG)13 |
| **GALK2** | ENSG00000156958 | ENST00000560138 | (TG)13 |
| **GALK2** | ENSG00000156958 | ENST00000559454 | (TG)13 |
| **RFX2** | ENSG00000087903 | ENST00000586806 | (T)12 |
| **RFX2** | ENSG00000087903 | ENST00000586806 | (T)12 |
| **RFC5** | ENSG00000111445 | ENST00000484086 | (T)12 |
| **PYCR1** | ENSG00000183010 | ENST00000582198 | (A)12 |
| **PYCR1** | ENSG00000183010 | ENST00000579366 | (A)12 |
| **DAGLB** | ENSG00000164535 | ENST00000436575 | (T)12 |
| **NPTN** | ENSG00000156642 | ENST00000565282 | (T)12 |
| **GLOD4** | ENSG00000167699 | ENST00000536578 | (A)12 |
| **EFHB** | ENSG00000163576 | ENST00000344838 | (T)12 |
| **ITGB1BP2** | ENSG00000147166 | ENST00000538820 | (T)12 |
| **KCNN4** | ENSG00000104783 | ENST00000615047 | (A)12 |
| **ACAT1** | ENSG00000075239 | ENST00000527942 | (T)12 |
| **MRPS36** | ENSG00000134056 | ENST00000512880 | (T)12 |
| **MRPS36** | ENSG00000134056 | ENST00000602380 | (T)12 |
| **MYO5C** | ENSG00000128833 | ENST00000558479 | (A)12 |
| **CHMP2B** | ENSG00000083937 | ENST00000472024 | (T)12 |
| **CHRFAM7A** | ENSG00000166664 | ENST00000299847 | (T)12 |
| **CHRFAM7A** | ENSG00000166664 | ENST00000562729 | (T)12 |
| **TTC14** | ENSG00000163728 | ENST00000492617 | (T)12 |
| **TTC14** | ENSG00000163728 | ENST00000495660 | (T)12 |
| **SLC6A9** | ENSG00000196517 | ENST00000475075 | (A)12 |
| **SNX1** | ENSG00000028528 | ENST00000560829 | (A)12 |
| **HS3ST4** | ENSG00000182601 | ENST00000331351 | (GCG)11 |
| **APOA2** | ENSG00000158874 | ENST00000468465 | (GT)11 |
| **APOA2** | ENSG00000158874 | ENST00000463812 | (GT)11 |
| **C1QB** | ENSG00000173369 | ENST00000510260 | (TGGA)11 |
| **C1QB** | ENSG00000173369 | ENST00000509305 | (TGGA)11 |
| **C1QB** | ENSG00000173369 | ENST00000432749 | (TGGA)11 |
| **RAPGEF5** | ENSG00000136237 | ENST00000458533 | (T)10 |
| **POLR3E** | ENSG00000058600 | ENST00000565358 | (T)10 |
| **PDXDC1** | ENSG00000179889 | ENST00000563522 | (T)10 |
| **PDXDC1** | ENSG00000179889 | ENST00000566426 | (T)10 |
| **PDXDC1** | ENSG00000179889 | ENST00000567306 | (T)10 |
| **PDE4DIP** | ENSG00000178104 | ENST00000479408 | (A)10 |
| **SRPK1** | ENSG00000096063 | ENST00000512445 | (T)10 |
| **TMBIM4** | ENSG00000155957 | ENST00000544599 | (T)10 |
| **DCTN4** | ENSG00000132912 | ENST00000424236 | (T)10 |
| **DCTN4** | ENSG00000132912 | ENST00000518015 | (T)10 |
| **DCTN4** | ENSG00000132912 | ENST00000521533 | (T)10 |
| **FZD8** | ENSG00000177283 | ENST00000374694 | (C)10 |
| **NPM1** | ENSG00000181163 | ENST00000521672 | (T)10 |
| **KITLG** | ENSG00000049130 | ENST00000552044 | (T)10 |
| **OR7A17** | ENSG00000185385 | ENST00000642123 | (AATA)10 |
| **OR7A17** | ENSG00000185385 | ENST00000641113 | (AATA)10 |
| **OR52N4** | ENSG00000181074 | ENST00000641350 | (T)10 |
| **UHRF1BP1L** | ENSG00000111647 | ENST00000545232 | (C)10 |
| **UHRF1BP1L** | ENSG00000111647 | ENST00000548045 | (C)10 |
| **UHRF1BP1L** | ENSG00000111647 | ENST00000551973 | (C)10 |
| **UHRF1BP1L** | ENSG00000111647 | ENST00000550544 | (C)10 |
| **UHRF1BP1L** | ENSG00000111647 | ENST00000551980 | (C)10 |
| **UGT2B7** | ENSG00000171234 | ENST00000502942 | (T)10 |
| **ABCF1** | ENSG00000204574 | ENST00000468958 | (A)10 |
| **ACAP2** | ENSG00000114331 | ENST00000439666 | (T)10 |
| **ARHGEF18** | ENSG00000104880 | ENST00000359920 | (T)10 |
| **ARL14** | ENSG00000179674 | ENST00000320767 | (A)10 |
| **ATXN3** | ENSG00000066427 | ENST00000502250 | (T)10 |
| **ATXN3** | ENSG00000066427 | ENST00000557311 | (T)10 |
| **EXTL3** | ENSG00000012232 | ENST00000523149 | (T)10 |
| **RAPGEF6** | ENSG00000158987 | ENST00000513227 | (T)10 |
| **NFIA** | ENSG00000162599 | ENST00000371191 | (T)10 |
| **ZNF91** | ENSG00000167232 | ENST00000595533 | (T)10 |
| **ZNF480** | ENSG00000198464 | ENST00000335090 | (T)10 |
| **TLR2** | ENSG00000137462 | ENST00000260010 | (GT)10 |
| **TSNAXIP1** | ENSG00000102904 | ENST00000388833 | (A)10 |

The human-specific STRs were non-existent in the orthologous genes at ≥3-repeats in 46 species studied across major classes of vertebrates (a total of 755,956 transcripts). The 5^th^ percentile of these genes is listed in Table 1 based on the length of the STRs (the entire list of the genes is presented as Suppl. 1). As an extreme example, the TIS of the *NVL* gene was flanked by a human-specific (T)22 STR, which is the longest STR detected in a human protein-coding gene TIS-flanking sequence. The TIS of the gene, *LRRC19*, was flanked by the longest poly-A at (A)20. Short and medium length STRs were also detected in the human-specific compartment (Suppl. 1).

Significant skewing was observed in the distribution of human-specific STRs vs. the overall (i.e. human-specific and non-specific) distribution of STRs in the human TIS-flanking sequence interval (Mann Whitney W=62532, p=1.4 × 10^−11^) (Fig. 2). For example, while the most abundant STR in the overall compartment was (T)6, the most abundant STR in the human-specific compartment was (CT)3. While the (GC)3 and (CG)3 dinucleotide STRs were enriched in the overall STR compartment, their abundance was significantly lower in the human-specific compartment. Instead, (CA)3 and (AC)3 were significantly more abundant in the human-specific compartment. Difference in the distribution of tri and tetranucleotide STRs was also observed between the two compartments. For example, while trinucleotide and tetranucleotide STRs of GC composition were more abundant in the overall compartment, non-GC STR compositions were more abundant in the human-specific compartment.

**Fig. 2.**
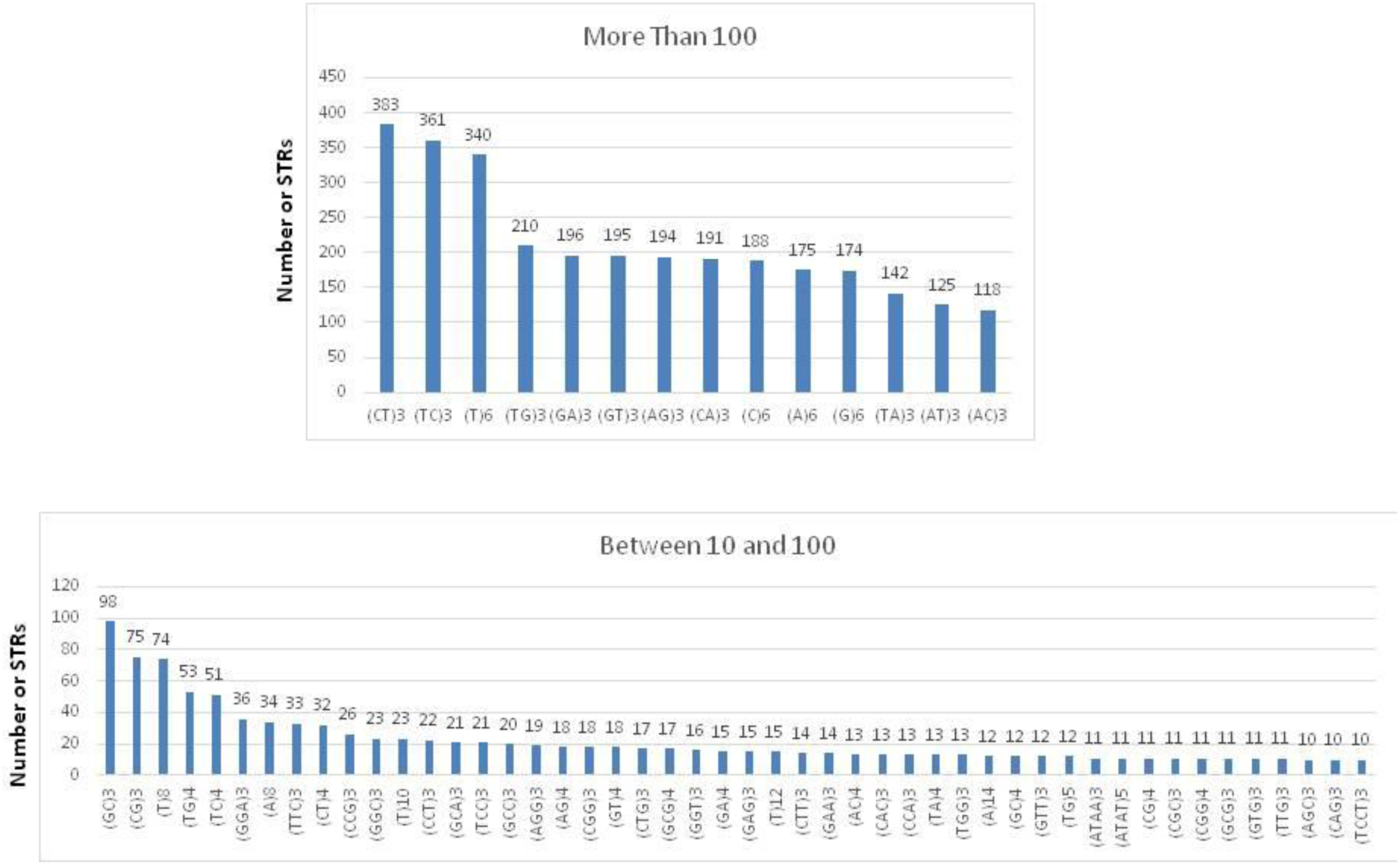
Distribution of the human-specific STR compartment in the TIS-flanking cDNA sequence. A significant skewing was observed between this compartment and the compartment containing the overall (human-specific and non-specific) STRs. The abundance of STRs is sorted in the ascending order.

### STRs link to TIS selection

We examined the hypothesis that there may be a link between STRs and TIS selection in human. The initial five amino acids (excluding the initial methionine) of the human protein sequences that were flanked by STRs were checked against all the initial orthologous five amino acids in 46 species across vertebrates in order to compare the number of events where human-specific and non-specific STRs occurred with homologous and non-homologous TISs (≥50% and <50% similarity of the five amino acids). Total of 2,025,817 pair-wise TIS comparisons were performed through the BLAST pipeline, and significant correlation was observed between STRs and TIS selection (p<0.00001) (Table 2), where there was a significant enrichment (2.98-fold) of non-homologous TISs co-occurring with human-specific STRs.

**Table 2.**
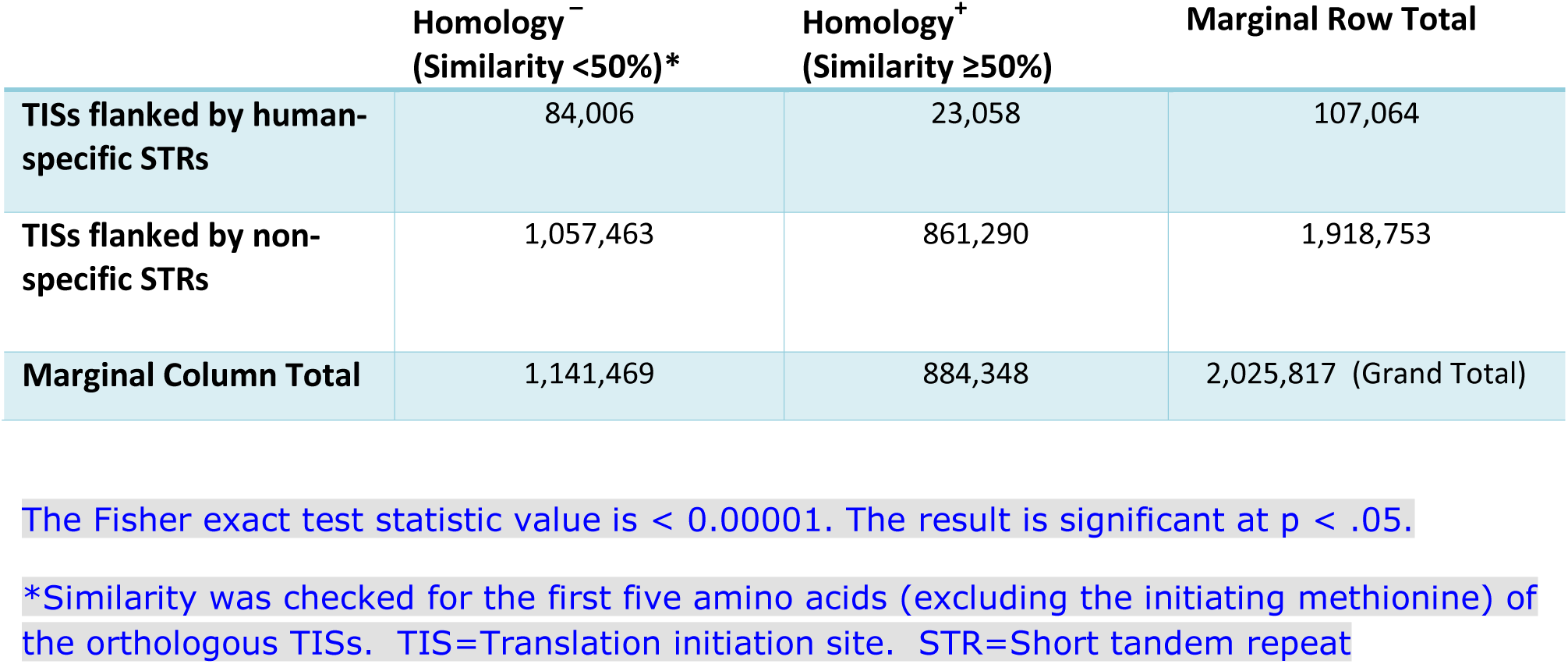
Evaluation of a link between STRs and TIS selection

| | Homology<sup>-</sup><br>(Similarity $< 50\%$)* | Homology<sup>+</sup><br>(Similarity $\geq 50\%$) | Marginal Row Total |
| --- | --- | --- | --- |
| **TISs flanked by human-specific STRs** | 84,006 | 23,058 | 107,064 |
| **TISs flanked by non-specific STRs** | 1,057,463 | 861,290 | 1,918,753 |
| **Marginal Column Total** | 1,141,469 | 884,348 | 2,025,817 (Grand Total) |

The Fisher exact test statistic value is $< 0.00001$ . The result is significant at $p < .05$ .
\*Similarity was checked for the first five amino acids (excluding the initiating methionine) of the orthologous TISs. TIS=Translation initiation site. STR=Short tandem repeat

## Discussion

The goal of this study was to characterize the STR landscape of the immediate 120 bp upstream sequence of human TISs at the whole-genome scale, to catalog the human-specific compartment of these STRs, and to investigate a possible link between STRs and TIS selection. Our findings provide the first indication of a link between STRs and TIS selection. The basis of this link was an excess of human-specific STRs co-occurring with non-homologous human TISs.

Sequence similarity searches can reliably identify “homologous” proteins or genes by detecting excess similarity (Pearson 2013). In our study, homology of the TISs was inferred based on three thousand random similarity scorings of the initial protein-coding five amino acids (excluding the initiating methionine), in which a similarity of ≥50% was considered as “homology”. This scoring methodology was consistently applied to the TISs linked to human-specific and non-specific STRs.

We also observed significant skewing of the human-specific STRs vs. the overall distribution of STRs. Genome-scale skewing of STRs, albeit at a lesser scale of STR classes, has been reported by our group in a preliminary study of the gene core promoter interval (Nazaripanah et al. 2018). The RNA structure influences recruitment of various RNA binding proteins, and determines alternative TISs (Martinez-Salas et al. 2013). Indeed, the ribosomal machinery has the potential to scan and use several open reading frames (ORFs) at a particular mRNA species (Kochetov et al. 2017). It is reasonable to envision that human-specific *cis* elements at the mRNA may result in the production of proteins that are specific to humans.

When located at the 5’ or 3’ UTR, STRs can modulate translation, the effect of which has biological and pathological implications (Rovozzo et al. 2016; Usdin 2008; Kumari et al. 2007). Remarkably, the disorders linked to the 5’UTR STRs encompass a number of human-specific neurological disorders. A number of the genes identified through our TIS-flanked STRs analysis (e.g. *NVL)*, confer risk for diseases that are predominantly specific to the human species, such as schizophrenia and bipolar disorder (Wang et al. 2015). Neurodegeneration is another example linked to genes such as *SULT1A3* (Butcher et al. 2017). In another remarkable example, *TBR1* is involved in *FOXP2* gene expression, which has pivotal role in speech and language in human (Becker et al. 2018). The above are a few examples of how the identified genes and their potential human-specific translation may be linked to human evolution and disease. Future studies are warranted to examine the implication of the identified STRs and genes at the inter-and intra-species levels.

## Conclusion

We present the landscape of STRs at the immediate upstream cDNA sequence flanking the human protein-coding gene TISs. Further, we found a link between STRs and TIS selection, which is supported by the skewing of the human-specific STR compartment. The data presented here have implications at the inter- and intraspecies levels and warrant further functional and evolutionary studies.

## Methods

### Data collection

Forty seven species encompassing major classes of vertebrates were selected, and in each species, the 120 bp upstream cDNA sequence flanking all annotated protein-coding TISs (n=755,956 transcripts) were analyzed based on the Ensembl database versions 90 and 91 (https://asia.ensembl.org). For each gene in each species, its Ensembl ID, the annotated transcript IDs, the coding DNA sequence (CDS) and the annotated cDNA sequences were retrieved (the list of species and investigated genes are available upon request). The CDS sequences and their annotated cDNAs were downloaded using REST API from the Ensembl database. The first start codon for each transcript was determined using BLAST between the CDS and cDNA. The 120 bp cDNA interval upstream of the start codon (ATG) was investigated for the presence of STRs (Fig. 3).

**Fig. 3.**
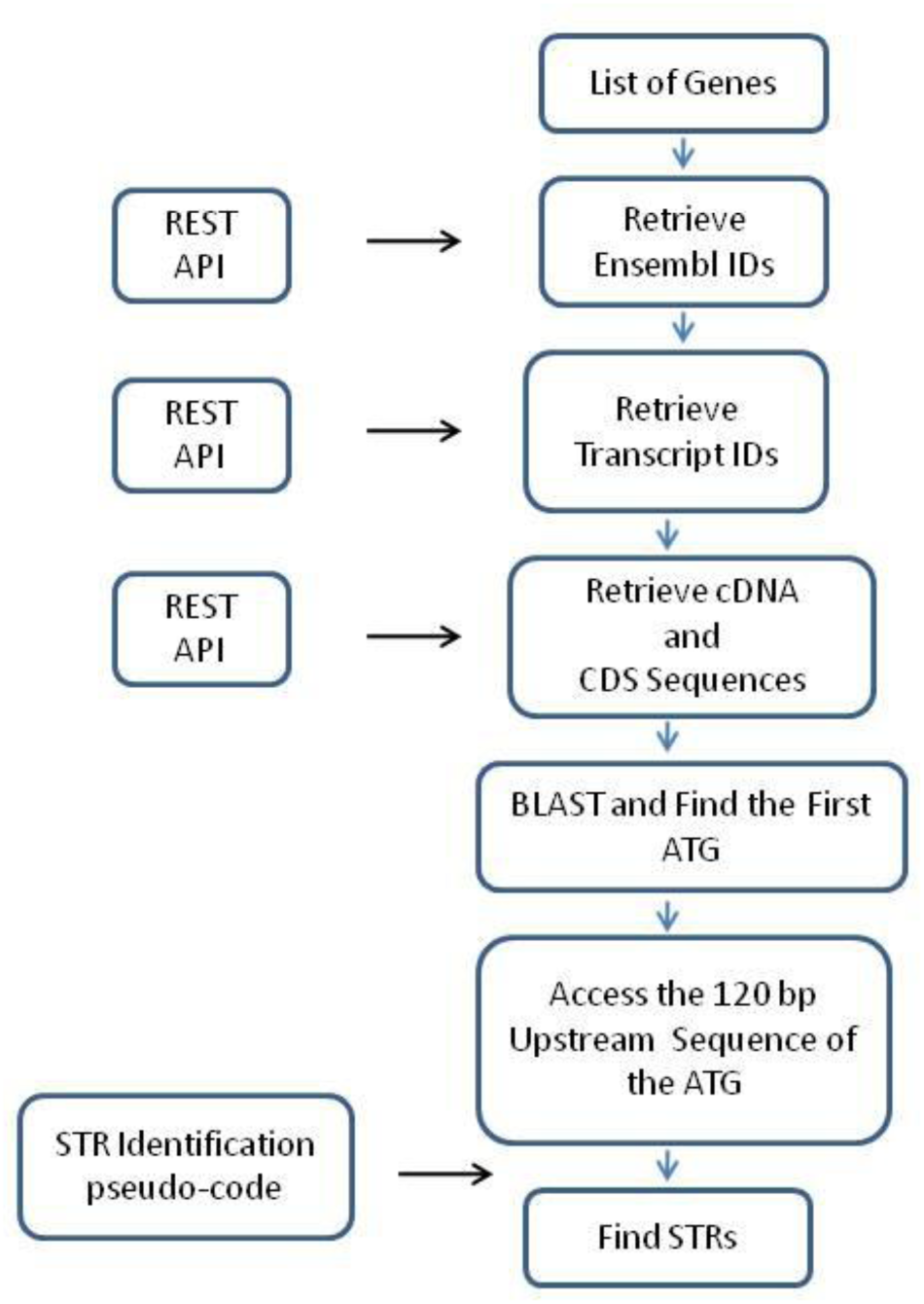
Workflow of STR identification in the TIS-flanking upstream sequence.

### Retrieval of gene IDs across species

Using the Enhanced REST API tools, a set of functions were developed to analyze genes and their transcripts information, including *func_get_ensemblID* and *func_get_TranscriptsID*. The cDNA and CDS sequences of genes and their respective transcripts were obtained using *func_get_GenomicSequence* and *func_get_CDSSequence* functions.

### Identification of STRs in the human TIS-flanking interval

A general method of finding human-specific and non-specific STRs for each individual gene was developed and applied as follows: The 120 bp cDNA sequence flanking the TIS of all annotated protein-coding gene transcripts was screened in 47 species across vertebrates for the presence of STRs. A list of all STRs and their abundance was prepared for each gene in every species. The data obtained on the human STRs was compared to those of other species and the STRs which were “specific” to human (i.e. not present at ≥3-repeats in any other species) were identified. The relevant pseudo-code for the identification of repeated substrings (STRs) is illustrated in Fig. 4.

**Fig.4.**
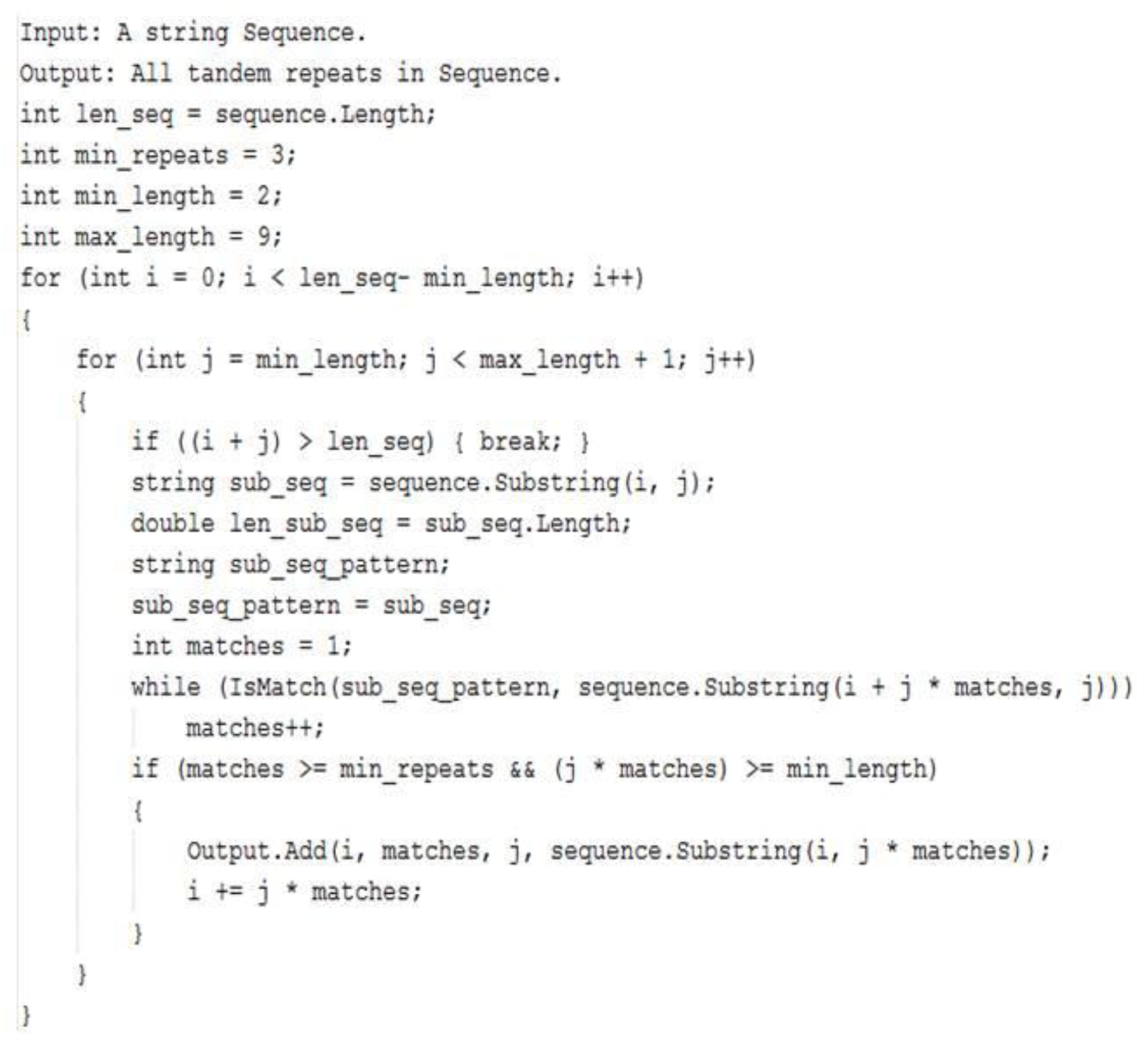
STR identification Pseudo-code

Mann-Whitney U test was used to compare the distribution of human-specific vs. the overall (specific and non-specific) STR distribution in human.

### Evaluation of a link between STRs and TIS selection

We hypothesized that there may be a link between STRs and TIS selection. Weighted homology scoring was performed, in which the initial five amino acid sequence (excluding the initial methionine) of the human TISs that were linked to STRs were BLASTed against all initial protein-coding five amino acids in 46 species across vertebrates (2,025,817 pair-wise TIS comparisons). The above was aimed at comparing the number of events where human-specific and non-specific STRs occurred with homologous and non-homologous TISs.

The following equation was developed for the weighted scoring of homology (Eq.1), where refers to the five amino acid sequence (excluding the initial methionine M) that are flanked by a STR at the cDNA sequence, *j* refers to the gene, and *B* refers to all the transcripts of the same gene that contain the STR in other species.

If M is the first methionine amino acid of two sequences, for all 5 successive positions represented by *i* in the equation, we defined 5 weight coefficients *W*_1_to *W*_5_ based on the importance of the amino acid position, observed in the *W* vector. The degree of homology between the two sequences *A* and *B* was calculated using function φ for all 5 positions with the operation (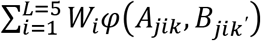) We repeated this operation *k* for transcripts, where *k* stands for the number of transcripts in human. *K’* refers to all transcripts of the gene *j* in other species.

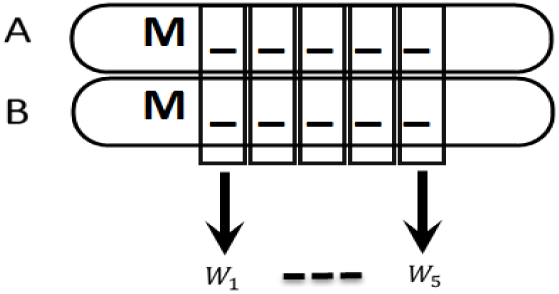

(1) 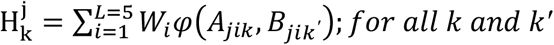

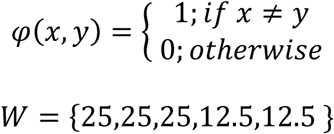

Homology of the five amino acids and, therefore, the TIS was inferred based on the %similarity scoring, in which a similarity of ≥50% was considered as “homology”. This threshold was achieved following BLASTing three thousand random pair-wise TIS five amino acids, which yielded a threshold of ≥50% (i.e. similarity scores ≥50% were considered “homology”). Finally, the two by two table and Fisher exact statistics were used to examine the link between STRs and TISs.

## Suppl. 1.

List of all human protein-coding genes which contain human-specific STRs in their TIS-flanking sequence.

